# Bayesian statistics improves biological interpretability of metabolomics data from human cohorts

**DOI:** 10.1101/2022.05.17.492312

**Authors:** Christopher Brydges, Xiaoyu Che, W. Ian Lipkin, Oliver Fiehn

**Author notes:** Corresponding author: Oliver Fiehn.

## Abstract

**Background:** Univariate analyses of metabolomics data currently follow a frequentist approach, using p-values to reject a null-hypothesis. However, the usability of *p*-values is plagued by many misconceptions and inherent pitfalls. We here propose the use of Bayesian statistics to quantify evidence supporting different hypotheses and discriminate between the null hypothesis versus lack of statistical power.

**Methods:** We use metabolomics data from three independent human cohorts that studied plasma signatures of subjects with myalgic encephalomyelitis / chronic fatigue syndrome (ME/CFS). Data are publicly available, covering 84-197 subjects in each study with 562-888 identified metabolites of which 777 were common between two studies, and 93 compounds reported in all three studies. By comparing results from classic multiple regression against Bayesian multiple regression we show how Bayesian statistics incorporates results from one study as ‘prior information’ into the next study, thereby improving the overall assessment of the likelihood of finding specific differences between plasma metabolite levels and disease outcomes in ME/CFS.

**Results:** Whereas using classic statistics and Benjamini-Hochberg FDR-corrections, study 1 detected 18 metabolic differences, study 2 detected no differences. Using Bayesian statistics on the same data, we found a high likelihood that 97 compounds were altered in concentration in study 2, after using the results of study 1 as prior distributions. These findings included lower levels of peroxisome-produced ether-lipids, higher levels of long chain, unsaturated triacylglycerides, and the presence of exposome compounds that are explained by difference in diet and medication between healthy subjects and ME/CFS patients. Although study 3 reported only 92 reported compounds in common with the other two studies, these major differences were confirmed. We also found that prostaglandin F2alpha, a lipid mediator of physiological relevance, was significantly reduced in ME/CFS patients across all three studies.

**Conclusions:** The use of Bayesian statistics led to biological conclusions from metabolomic data that were not found through the frequentist analytical approaches more commonly employed. We propose that Bayesian statistics to be highly useful for studies with similar research designs if similar metabolomic assays are used.

## Introduction

Although literature detailing limitations and misconceptions of *p*-values has established that the way that many researchers employ statistics in their research is ritualistic and inappropriate,^1,2^ and impedes scientific progress,^3,4^ metabolomic researchers rarely acknowledge the pitfalls in methods used for assessment of statistical significance. Even the American Statistical Association has warned about misusing the p-value.^5^ Cut-offs for significance testing can be easily subject to changing perspectives.^6-8^

This discussion has not become generally accepted in the realms of omics sciences, including in metabolomics. While it is common for univariate analyses to be conducted with false discovery rate or family-wise error rate corrections, researchers still only consider metabolites to be interesting if their *p*-values fall below an arbitrary threshold. But *p*-values only give the probability that measured data fit an assumed null hypothesis. If a *p*-value is <0.05, the null hypothesis is commonly rejected. However, even if *p*>0.05, a metabolite might still be different between two study groups. Simply put, strict “null hypothesis significance testing” cannot distinguish whether there is a true null effect or whether the data are insensitive.^9^ More importantly, a *p*-value cannot provide support for an alternative hypothesis. A classic *p*-value reports the probability of the data given the hypothesis – not the probability of the hypothesis, given the data.^5^ These are not identical statements (i.e., p(data|H) ≠ p(H|data)),^10^ and they do not answer the same questions. Strangely, a *p*-value does not provide a measure of the strength of evidence in favor of a hypothesis. One cannot rank *p*-values by being ‘more significant’. For example, a *p*-value of 0.0001 is not stronger evidence in favor of the alternative hypothesis than a *p*-value of 0.049. Given a threshold of 0.05, they both reject the null hypothesis, but neither of them tests an alternative hypothesis. Furthermore, although reporting actual effect sizes (the differences in metabolite levels) is crucially important, a *p*-value does not give that information.

We here present Bayesian alternatives to complement or replace *p*-values and the analyses that produce them that we will hereafter refer to as null hypothesis significance testing (NHST). We exemplify the power of Bayesian analyses on previously published metabolomics data to highlight the differences between the two statistical approaches. But how is Bayesian statistics different? Unlike classic *p*-values, Bayesian statistics can be used to quantify the size of an effect, quantify the strength of evidence in favor of one hypothesis over another, and allow researchers to discriminate between an inconclusive finding and evidence in favor of the null hypothesis. We argue that this is exactly what researchers want to get. These analyses test competing models of hypotheses and the distribution of the observed data.^11,12^ Bayesian statistics always starts with a prior distribution. That is an expectation of a range of possible effect sizes that could feasibly be observed. Put simply, this means that researchers should (or at least can) have some idea of the plausible size and/or direction of an effect that they are studying. For example, a statement like “I think that compound X is most likely to be upregulated in cases compared to controls, with a fold-change of about 3” is an example of a prior distribution: this description can be modeled very easily so that the probability of downregulation is zero, and the most likely effect sizes are around fold change = 3.

Bayesian statistics are used across a vast array of fields and domains,^13^ including everyday cognition and decision-making.^14^ An example is hurricane forecasts. **Figure 1** shows a fictional hurricane that may have formed in the Gulf of Mexico (top-left panel), with a cone of different colors displaying the path the hurricane is most likely to take. Given previous hurricanes, and other data such as ocean currents, storm trackers already have an expectation, a “prior distribution”. This is like a metabolomicist knowing the literature and having ideas how levels of metabolites might vary in diabetes mellitus. The colors of the cone correspond to the modeled prior distribution (bottom-left panel). As the hurricane progresses (Figure 1, top-right panel) and researchers incorporate the new data into an updated model, their expectations change about where the hurricane is most likely to go. The cone also narrows as the researchers become more confident about the path of the hurricane. This results in the narrower distribution (bottom-right panel). Hence, unlike classic *p*-values, Bayesian models take prior data into account and can be updated continuously.

**Figure 1.**
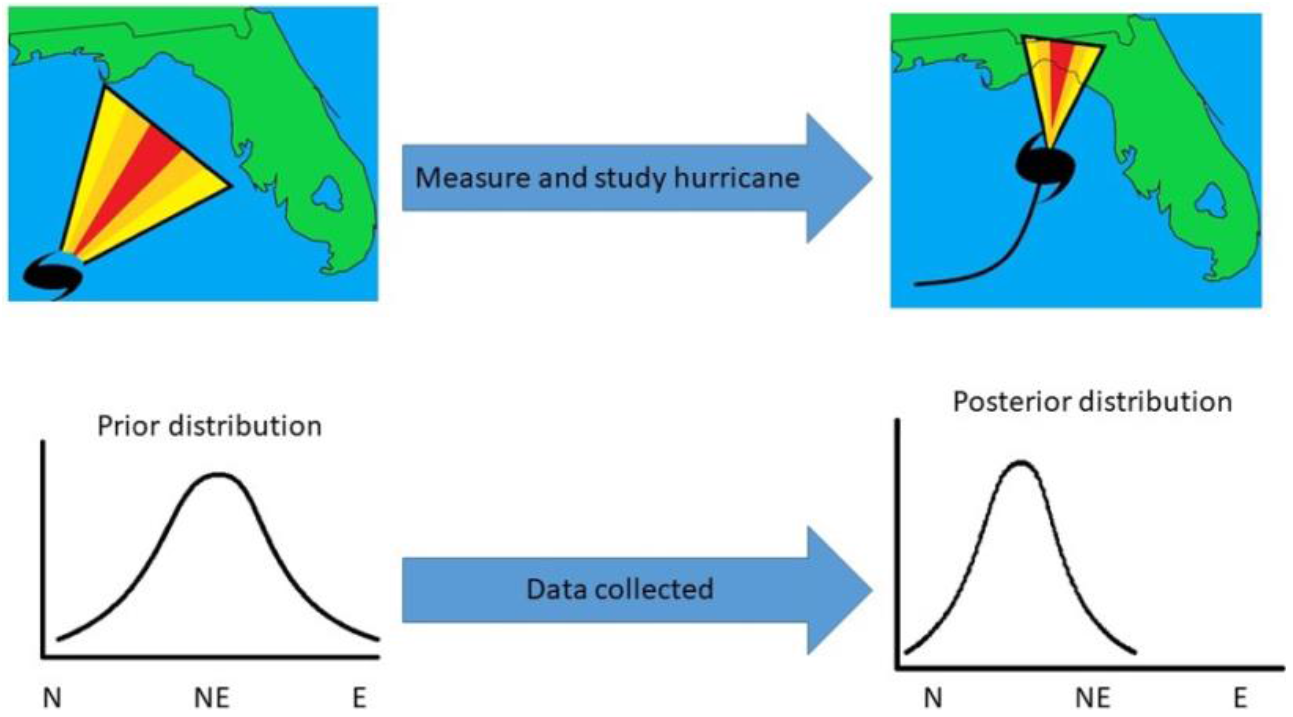
Hypothetical hurricane paths before (top-left) and after data has been collected (top-right). The cones represent the likely path of the hurricane (yellow = unlikely; red = most likely). Distributions (bottom-left and bottom-right) represent the cones plotted as probability distributions.

## Methods

### Metabolomic Datasets

Three data sets were reanalyzed for the current study. All three studies investigated metabolic differences between patients who suffer from myalgic encephalomyelitis/chronic fatigue syndrome (ME/CFS), and matched healthy controls. In order to be consistent in our analyses, we only used identified metabolites and ignored unknown features. First, the previously published study by Nagy-Szakal et al^15^ included 50 cases and 50 controls and reported 562 identified metabolites. The data is available at the Metabolomics Workbench repository^16^ under Project ID PR000576 (DOI: http://dx.doi.org/10.21228/M86X1F). The second data set is available under Project DOI 10.21228/M8PD9N and available by Che et al^17^ at DOI: 10.1101/2021.06.14.21258895, covering 106 cases and 91 control subjects, and reporting 888 metabolites. In both studies, participant metadata (age, sex, body mass index (BMI), race/ethnicity, diagnosis of irritable bowel syndrome (IBS), geographic/clinical site, and season of sample) was also collected. 20 subjects (10 ME/CFS patients, 10 healthy controls) participated in both studies. For comparison of the strengths of Bayesian analyses, we also used a previously published data set by Naviaux et al^18^ (Project DOI: 10.21228/M82K58) with 45 cases and 39 controls. Naviaux et al. used a different metabolomic assay with 612 identified metabolites, for which we found only 92 compounds in common to the other two studies. For this study, participant metadata as given above were not available apart from disease status and sex.

### Statistical Analyses

All analyses were conducted in R 4.1.2. Prior to analyses, any compound not observed in at least 50% of samples was removed from analysis. Any missing data were imputed with half-minimum values. Each compound was log transformed for normality and then auto-scaled. Only compounds common to both *Nagy-Szakal* and *Che* data sets were used in analyses, which resulted in 632 compounds (551 identified) being analyzed. Common compounds were matched using International Chemical Identifier keys. For the third, *Naviaux* data set, 92 compounds were matched to the other two data sets, using RefMet annotations^19^. Classic univariate statistical analyses were performed by linear regression using base R functions, and Bayesian regression was conducted using the rstanarm^20^ and bayestestR^21^ packages. These analyses were used to determine between-groups differences for each compound and were used instead of *t*-tests or Mann-Whitney *U* tests due to the inclusion of covariates and having a sufficiently large sample size. All models included age, sex, BMI, race/ethnicity, IBS diagnosis, geographic/clinical site, and season of sample as covariates. The default prior distributions recommended by rstanarm were used to model the expected effect sizes for each compound in the *Nagy-Szakal* data set. These defaults are considered to be ‘weakly informative’ in that they provide some information on the expected magnitude of the effect based on the scales of the variables. However, they do not strongly affect the posterior distribution and help stabilize computation, while still allowing for extreme effect sizes if warranted by the data.^22,23^ Posterior distributions were created from four Markov Chain Monte Carlo chains of 2,000 iterations each, with the first 1,000 iterations in each chain used as burn-ins. The posterior distribution of each compound from the *Nagy-Szakal* data was then used as the prior distribution for the same compound in the *Che* data set. Compounds were considered to be altered if the 95% credible interval did not overlap with zero. A negative posterior median was indicative of downregulation in the ME/CFS group, and a positive posterior median was indicative of upregulation. ChemRICH^24^ was performed for set enrichment statistics, with the posterior median used as an estimate of effect size and the probability of direction was used as a Bayesian analog of a *p*-value.^25^

## Results

### Classic univariate statistics analyses

Figure 2. shows the results of the classic *p*-value (frequentist) analyses for the *Nagy-Szakal* and *Che* data sets. Of the 632 compounds common to both studies, only 18 were significantly different (FDR < 0.10) in the *Nagy-Szakal* data for which phosphatidylcholines (28:0, 30:0, 32:1, 32:2, 33:0, 34:1, 34:3, 34:4, 38:2, 38:6) were found downregulated and triacylgycerides (52:4, 54:6, 54:7, 56:5, 56:8) were upregulated, in addition to lower levels of carnitine and tyrosine in the ME/CFS group. Interestingly, when analyzing the *Che* data set by itself, not a single metabolite was found to be significantly different between groups, using regression analyses (**Figure 2**).

### Bayesian Analyses

To demonstrate the advantages of Bayesian analyses over classic univariate analyses, we first analyzed the *Nagy-Szakal* data using weakly informative prior distributions. The resulting posterior distribution was then used as input into the Bayesian analyses of the *Che* data. Bayesian model comparison commonly refers to the calculation of Bayes factors (BFs). Bayes factors are ratios that quantify the probability of one hypothesis over another by estimating the strength of evidence.^26^ Such tests do not determine if one hypothesis is true and the other is not. but instead, whether one hypothesis is more likely than an alternative hypothesis, given the observed data. Additionally, BF values are easily interpretable in terms of the strength of a finding. Bayesian analyses allow statements such as “the null hypothesis was five times more likely to be true than the alternative hypothesis, given the data”. Classic *p*-values cannot support such statements. BFs are continuous estimates that range from approaching zero to approaching infinity. BF=1 indicates equal likelihoods of either hypothesis, given the data. Values further from 1 imply stronger evidence in favor of one hypothesis over the other. Jeffreys^27^ provided arbitrary guidelines to categorize these values as anecdotal, moderate, strong, or extreme evidence for or against one hypothesis over another. Based on these guidelines, BFs between ⅓ and 3 are referred to as anecdotal evidence and imply that the data are not sensitive enough to conclusively state that one hypothesis is more likely than another. For studies with BFs between are ⅓ and 3 are typically underpowered, and more data would need to be collected. However, any general rule used to categorize BFs will not be appropriate for all research contexts. Extraordinary claims require extraordinary evidence: if geographic location or birth date would have been found to be associated with ME/CFS, one would certainly require more evidence than usual for such claim.

**Figure 2.**
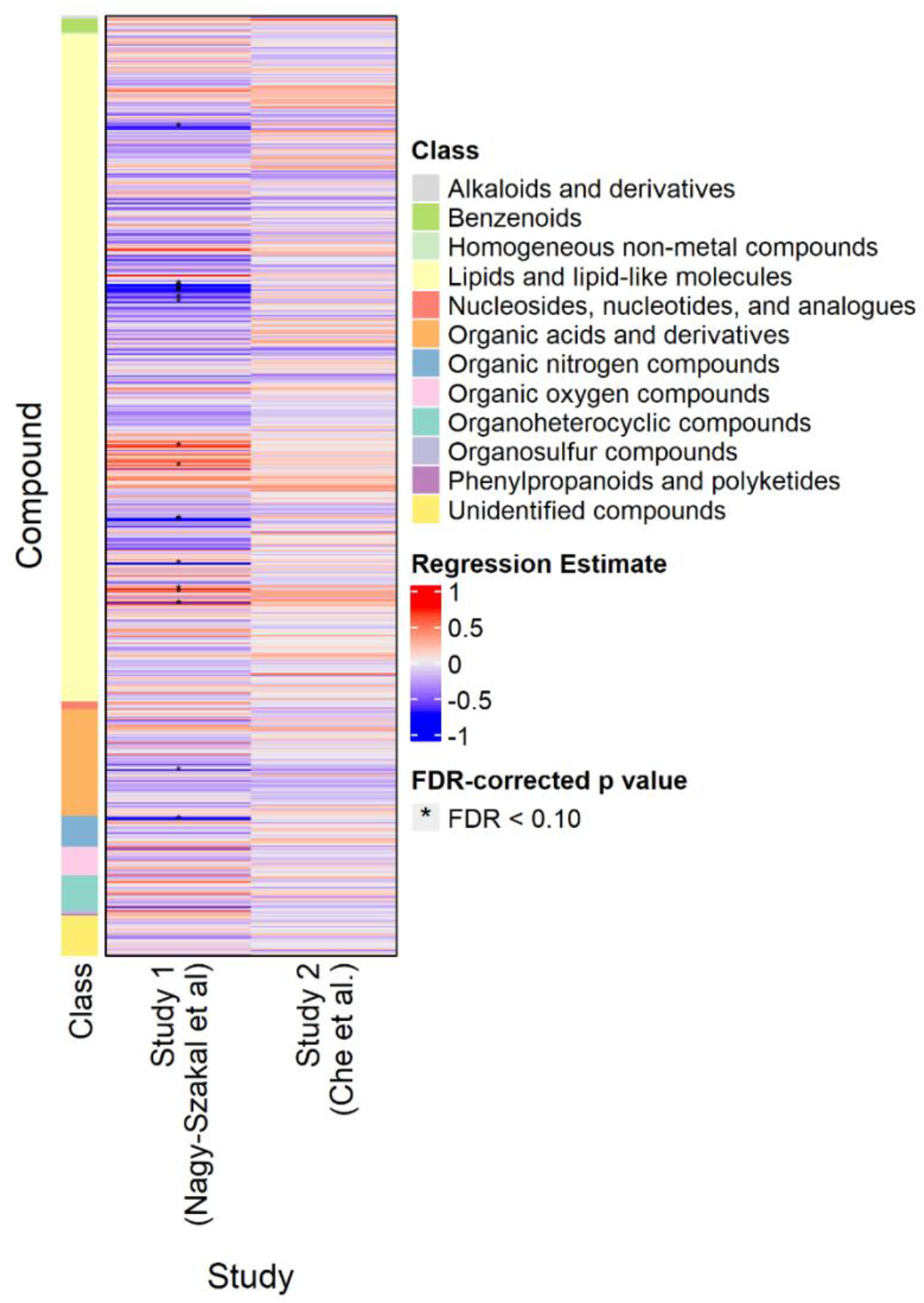
Heat map of compounds analyzed in the *Nagy-Szakal* and *Che* data sets. Blue indicates downregulation, red indicates upregulation. Asterisks indicate FDR-adjusted *p* values < 0.10.

Compounds with BFs> 10 closely match those that were reported as significantly different in the classic univariate analyses, for the simple reason that both methods tested whether metabolite levels were different between the case and control groups. For example, phosphatidylcholines (PC) were still found downregulated by Bayesian statistics, and triacylglyceride (TG) 54:7 was still upregulated in ME/CFS patients (**Figure 3**). For compounds with BF > 3, see Supplement 1. However, unlike the classic statistics, Bayes’ models also determine which compounds are sufficiently unlikely to be altered by ME/CFS (those with a BF < 1/3) and those that may be affected but do not have sufficient statistical power to draw a conclusion (BFs between 1/3 and 3).

**Figure 3.**
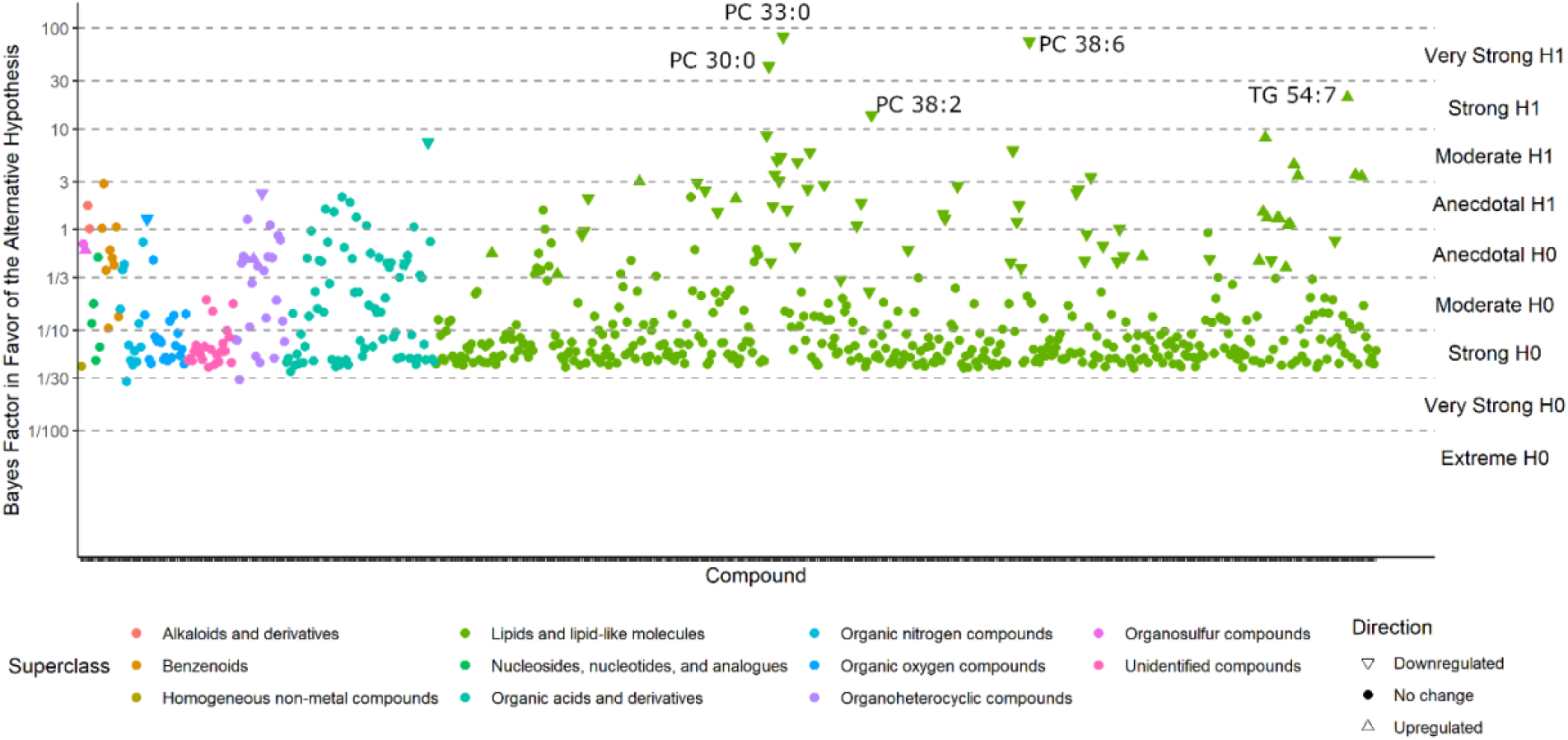
Bayes factors of all compounds in the *Nagy-Szakal* data set. Compounds with BF > 10 (i.e., strong evidence in favor of the alternative hypothesis) have been labeled.

In addition, Bayes’ analyses can be used to estimate the magnitude of an effect of interest. It is based on the posterior distribution, the updated prior distribution after data has been collected and incorporated into the model. The posterior distribution provides an estimate of the credibility of every possible metabolite value after accounting for the researcher’s hypotheses before data collection and after observing the current data. The mode of the distribution is the most likely estimate of the true effect size, while the mean and median of the distribution are often used as point estimates of credibility. The variability of a Bayesian effect size estimate is based on ‘credible intervals’. Credible intervals are a range of values within which the true effect falls at a specified level of confidence.^27^ When a 95% credible interval does not overlap with zero, the probability of an effect being zero is < 5%. Conversely, if the credible interval would overlap with zero, it could be considered as a Bayesian analog of a significance test.^25^ Notably, the similar-sounding ‘confidence intervals’ are different because those values would provide an interval within which the true effect size would fall at 95% confidence if the same study was to be repeated 100 times.^3^

Figure 4. shows the compounds that have 95% credible intervals that do not overlap with zero and have a BF > 3. Again, phosphatidylcholines were found downregulated along with tyrosine and one phosphatidylethanolamine (PE) ether-lipid, PE (p-38:6). A range of triacylglyerides were found up-regulated in ME/CFS cases, as also observed with classic univariate statistics. The most interesting difference in using Bayesian statistics was found when then using the *Che* data set with a prior distribution for each compound taken from the results of the *Nagy-Szakal* data set as. **Figure 5** shows the BFs of each compound, and **Figure 6** shows the compounds that have 95% credible intervals that do not overlap with zero. In contrast to classic univariate *p*-values (**Figure 2**), 98 compounds were found to be altered in the Bayesian analyses. Importantly, results were very consistent with those observed from the *Nagy-Szakal* data set: specific phosphatidylcholines were still found downregulated whereas specific triacylglyerides upregulated. Yet, a range of other compounds were now found differentially regulated as well: With Bayesian statistics informed by prior research (here: the *Nagy-Szakal* data), the *Che* data now showed that the branched chain amino acid leucine and aromatic amino acids tyrosine and phenylalanine were downregulated in ME/CFS cases. Additional lipid species were now also found to be down-regulated such as specific lysophosphatidylcholines, phosphatidylcholines, and plasmalogens. Notably, specific diacylglycerides were found at higher plasma levels in ME/CFS patients as well as specific pharmaceutical drugs such as gabapentin and p-acetamidophenol (acetaminophen), both used as pain medications, in addition to pantothenic acid (vitamin B5), a vitamin often taken as dietary supplement. A range of food compounds were found at lower levels in ME/CFS patients indicating less use of specific foods, such as caffeine, theobromine and trigonelline (coffee biomarkers) and piperine (found in pepper).

**Figure 4.**
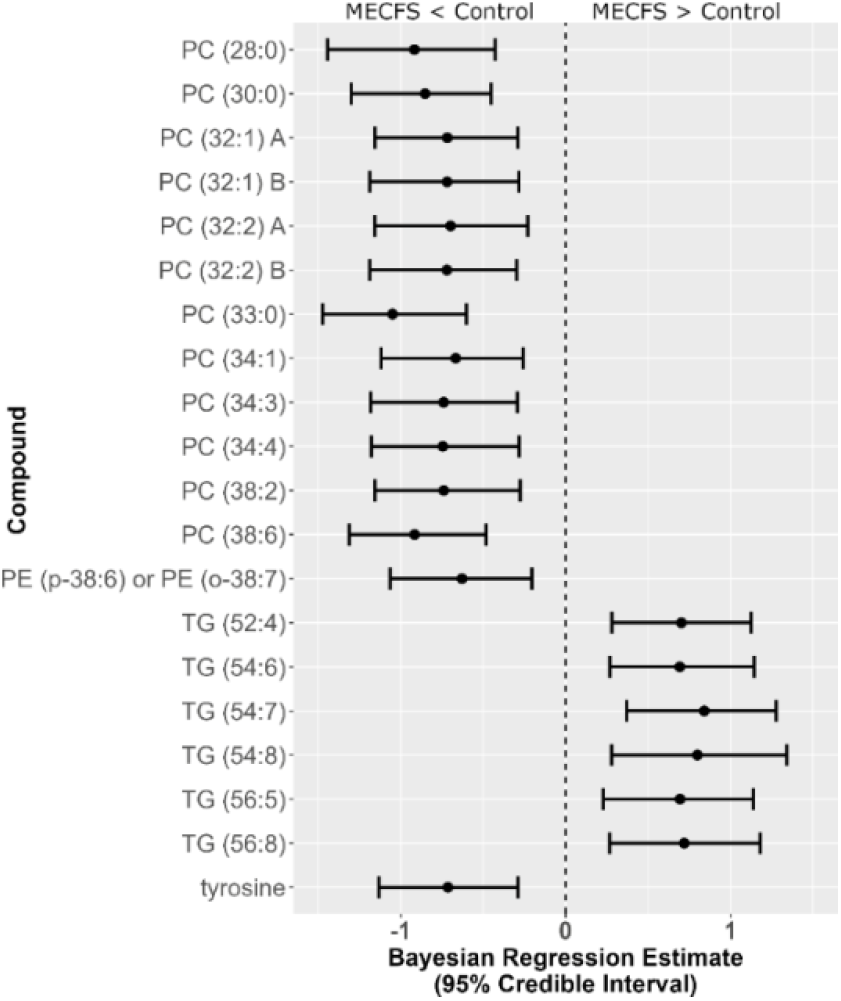
Forest plot of compounds with Bayesian 95% credible intervals not overlapping with zero and BF > 3 in the *Nagy-Szakal* data set. Points in each bar represent posterior distribution median and bars represent 95% credible interval bounds.

**Figure 5.**
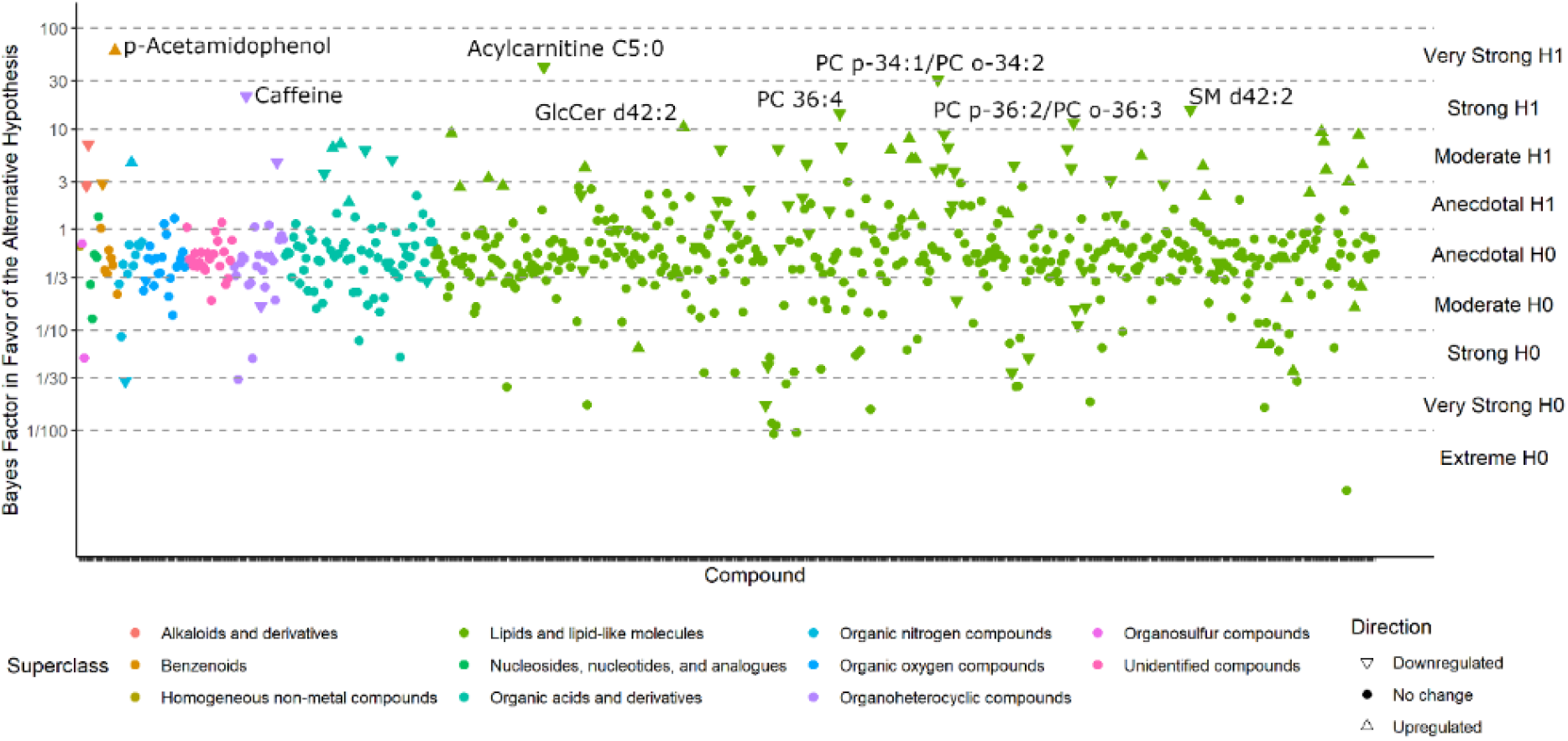
Bayes factors of all compounds in the *Che* data set. Compounds with BF > 10 (i.e., strong evidence in favor of the alternative hypothesis) have been labeled.

**Figure 6.**
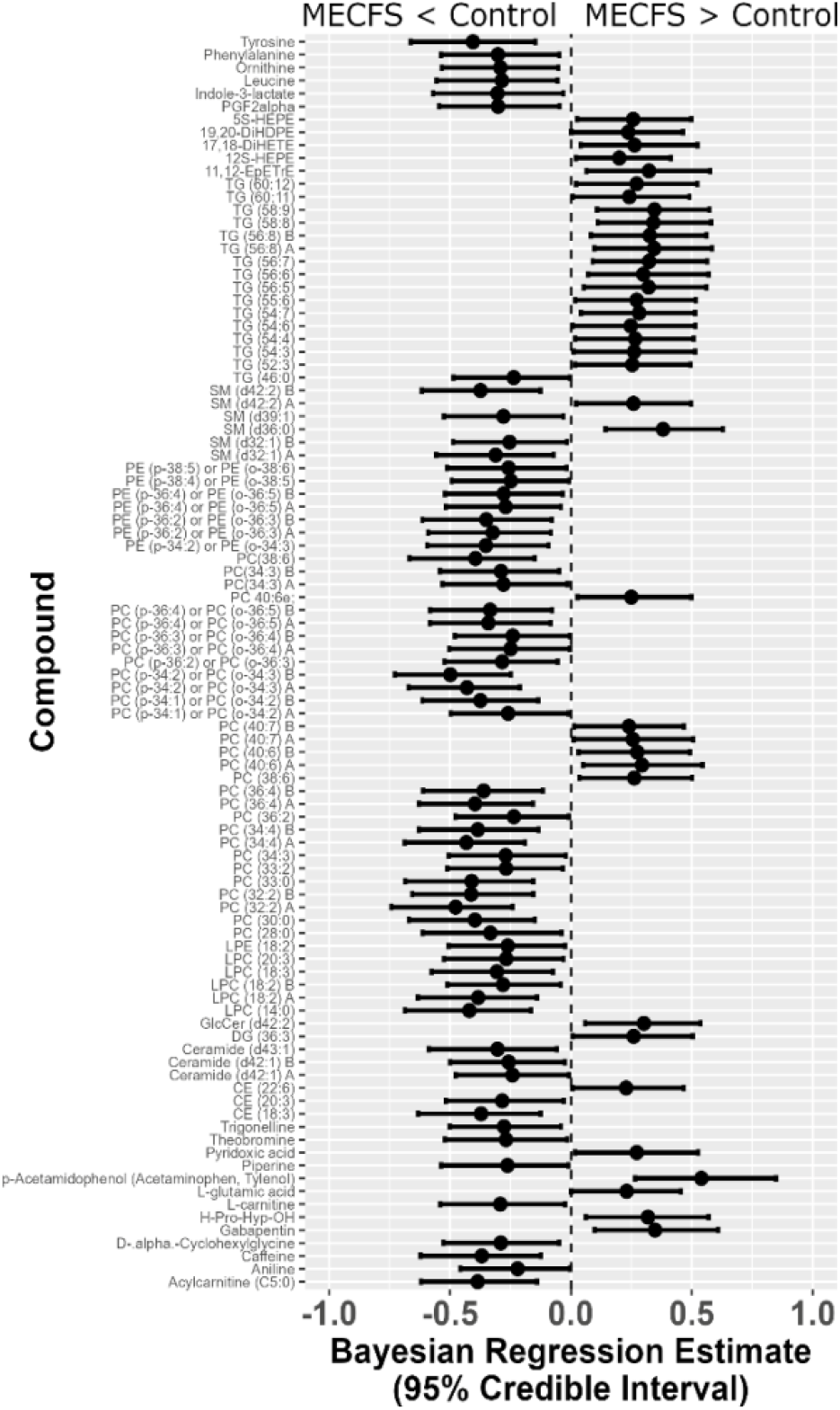
Forest plot of compounds with Bayesian 95% credible intervals not overlapping with zero in the *Che* data set. Points in each bar represent posterior distribution median and bars represent 95% credible interval bounds

There is one more type of question that metabolomics investigators, and their clinical partners, would ask: how are these findings connected? One way to answer such questions is to map metabolites to their biochemical pathways of synthesis and degradation. However, metabolite levels in human blood do not only correspond to pathways in cells but surely also to the differences between organs, and, of course, dietary patterns and exposures. We therefore tested for the significance that specific groups of compounds were found to be over-enriched, beyond what would be expected at random. For this type of set enrichment statistics,^29,30^ we have grouped metabolites by similarity in chemical structure, using the Kolmogorov-Smirnov test for significance. Using the results from Bayesian probabilities as input into chemical enrichment statistics,^24^ we found very strong evidence of very specific differential regulation of whole groups of compounds (**Figure 7**). For example, we found that only unsaturated triacylglycerides were upregulated in ME/CFS patients, but not saturated triacylglycerides. Such finding points to a specific biochemical regulation instead of simple explanations like differences in the number or type of lipid-carrying lipoprotein particles. Similarly, we found large downregulation of unsaturated phospholipid ethers and plasmalogens that are exclusively produced by peroxisomes and which, hence, might be involved in the etiology of the disease. Sphingomyelins and unsaturated phosphatidycholines were significantly associated with ME/CFS, but among these compound classes, some members were found to be up-regulated and others down-regulated. Unsaturated ceramides and unsaturated lysophosphatidylcholines were found down-regulated, but not their saturated counterparts. In combination, therefore Bayesian analyses unequivocally found evidence that reflects know behaviors in ME/CFS patients (such as avoidance of specific foods but increases in use of pharmaceuticals), in addition to biochemically and physiologically interpretable findings that would have been completely overlooked and ignored by classic univariate statistics.

**Figure 7.**
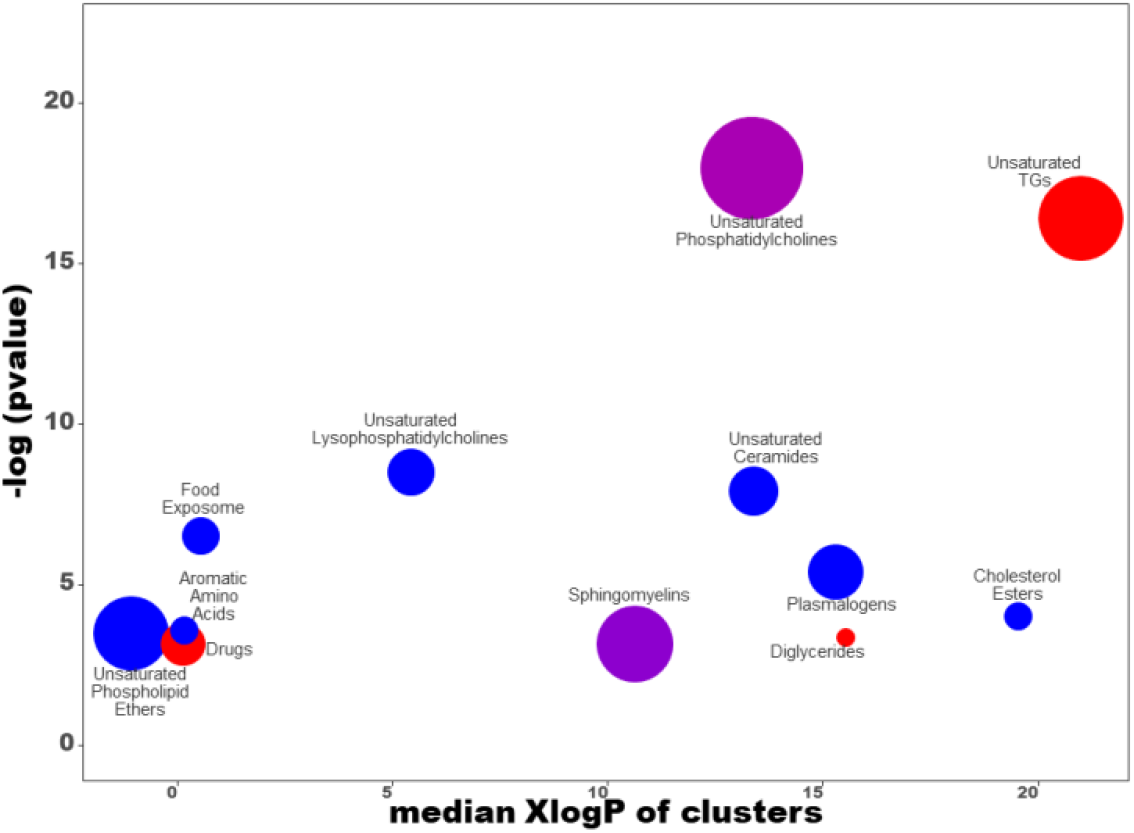
Chemical set enrichment statistics using results of the Bayesian analyses conducted on the *Che* data set. Bubble sizes indicate the number of compounds belonging to chemical groups. Bubble colors indicate the direction of effects (red = all compounds upregulated in ME/CFS group; blue = all compounds downregulated in ME/CFS group; purple = mixed effects).

Yet, Bayesian statistics can go even further. To illustrate the usefulness of Bayesian analyses, we extended our investigations by taking a third ME/CFS data set into account, the *Naviaux* study, **Figure 8**. Compounds were matched across all three data sets using RefMet,^19^ yielding a common core of 92 analyzed compounds that were observed in all three studies. The posterior distributions from the *Che* data were used as the prior distributions in the *Naviaux* study. Bayesian statistics yielded credible intervals for 14 compounds to be different between ME/CFS subjects and that were common for all three studies. Similar to the combined Bayes analysis of just the *Nagy-Szagal* and *Che* studies (Figure 6), we found specific phosphatidylcholines to be downregulated along with aromatic amino acids phenylalanine and tyrosine, while vitamin levels were elevated in ME/CFS patients (pantothenic acid and 4-pyridoxic acid). Interestingly, eicosapentaenoic acid (an omega-3 fatty acid and precursor to oxylipins) and PGF2-α (an oxylipin product and physiologically active mediator) were found to be regulated in opposite directions, further positing interesting new biochemical hypotheses on the etiology of ME/CFS that are now grounded in three independent data sets. Results like these also show the need to further standardize metabolomics data acquisitions that would accomplish progress in direct data integrations by Bayesian statistics, as demonstrated for this ME/CFS data integration study.

**Figure 8.**
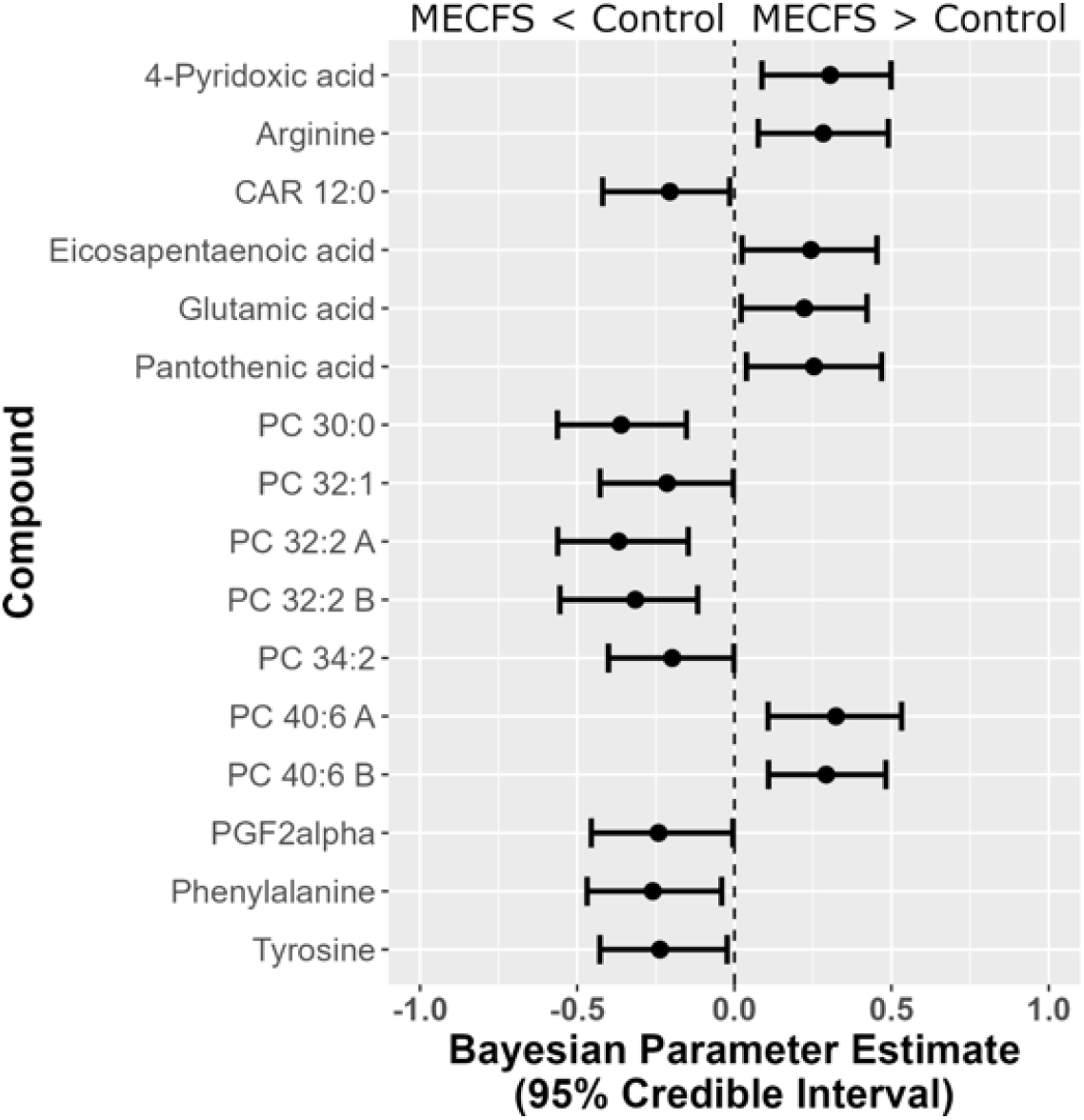
Forest plot of compounds with Bayesian 95% credible intervals not overlapping with zero in the *Naviaux* data set. Points in each bar represent posterior distribution median and bars represent 95% credible interval bounds.

## Discussion

Science should progress over time in its insights into specific phenomena. Bayesian statistics allow researchers to incorporate previous knowledge into their models. Hence, it is surprising that Bayesian approaches are rarely used in –omics research, and even less in metabolomics. Here, we exemplified the suitability of Bayesian approaches on three data sets that investigate the metabolic profile of ME/CFS. ^15,17,18^ There is little known about ME/CFS with respect to origins of this complex disease, disease progression, possible treatment options, chances or timing of remission or why women are more affected than men. ^31^ These studies were chosen precisely because the overall effects are not well studied, and because they investigated the same disease complex. The similarity in the study designs, including similar numbers of subjects, allowed us to conduct Bayesian analyses on the first data set, then incorporate the knowledge gleaned from those results into the Bayesian analyses conducted on the second and then the third data set. However, as caveat, we must state that there are likely differences in the ME/CFS cohorts that are unknown to us or to the different study investigators, because the disease etiology cannot be very sharply defined. It is very possible that the disease symptoms that are today summarized as ME/CFS may be better categorized into subtypes in the future. Hence, by random chance, there may have been differences in the ME/CFS subjects (or the matched healthy controls) that could have led to differences in metabolic imprints in the plasma metabolome data studied here. While all three studies used here emphasized the importance of complex lipids in plasma metabolomic profiles, the use of Bayesian analyses allowed us to refine the original ideas and metabolic pathways involved. For example, Naviaux et al. emphasized the importance of sphingolipids as having the ‘largest disturbances in the chemical signature of CFS’,^18^ explaining 44-50% of the metabolic impact in men and women.^18^ With the results by Nagy-Szagal^15^ and Che^17^, published later, we can now rule out these molecules as being of high importance when integrating the studies. Instead, we were able to use prior data such as the ceramides and sphingomyelins as hypotheses to be tested, but changing distributions and therefore altering posterior distributions, effect size estimates and the strengths of statements using subsequent datasets. Indeed, using Bayesian statistics and incorporating previous results into subsequent analyses we found evidence of peroxisomal dysfunction, differences in diet, and PGF2 alpha in the *Che* data set that we would not have otherwise seen if we only used traditional statistical analyses. Similar to other statistical approaches, Bayesian analyses can only weigh between different hypotheses but cannot ultimately provide absolute statements. For example, as Bayesian analyses provide strong evidence of the involvement of peroxisomes in the pathology of ME/CFS, we cannot rule out that mitochondria or the endoplasmic reticulum are also involved through the oxidation and modification of complex lipids. Similarly, our analyses focused on patient plasma which almost always precludes definitive conclusions on the timing and involvement of specific organs. Future research may use animal models and possible timing of events to study routes from potential initial causes (such as viral infections that lead to immune hyper-responses) to secondary defects (such as peroxisomal damages in liver and brain) which then may end in differences of physiologically active lipid mediators such as PGF2alpha that may lead to lower blood flow to brain regions, causing pain and brain fog as reported by ME/CFS patients.

Although the analyzed data sets we used are from metabolomics backgrounds, such tests can be applied to any field of quantitative hypothesis-driven research where classic univariate (p-value driven) analyses are traditionally used. Statistical analyses of metabolomics data have typically followed a predefined routine of conducting a series of univariate tests, such as *t*-tests/ANOVAs or their non-parametric equivalents.^32-34^ For effect size considerations, the current practice is even more dismal: fold changes are often (but not always) reported, but even these are limited as they do not take the within-compound variation into account.^35^ Bayesian analyses are advantageous as they can consider both the likelihood of a hypothesis being true and an estimation of effect size at the same time. Researchers should consider reporting *p*-values, effect sizes – whether fold changes or a standardized measure of effect size like Cohen’s *d* – and Bayesian results so that readers can gain greater insight from the data.^36^ Entirely Bayesian analyses, which report Bayes Factors, posterior estimates, and credible intervals, may also be suitable when designing a study.

Although Bayesian model comparisons can be conducted on any set of competing models,^37^ we here focused on simple case/control-style analyses. More advanced Bayesian analyses can be employed to answer increasingly complex and specific research questions, and do not have to be restricted to linear relationships.^38^ Researchers who add these analytical skills to their statistical toolbox will be able to elicit richer, more detailed information from their data than others who simply report a *p*-value.

There are some limitations of Bayesian statistics. First, the potentially subjective nature of selecting a prior distribution is a common criticism,^37^ as researchers may attempt multiple analyses in order to find a favored result using different prior distribution parameters. Therefore, researchers should aim to be transparent with their choice and justification for their priors and their statistical analyses. Choosing a suitable prior distribution can also be challenging, particularly for researchers unfamiliar with Bayesian analyses.^28^ Researchers should consider the following issues when deciding upon a prior distribution: The expected effect size and its potential variability, whether the hypothesis is one- or two-tailed, and the researcher’s confidence in observing a relatively specific effect size.^39^ Limitations of BFs also include their interpretation. BFs only provide relative, and not absolute, evidence for a hypothesis: a statement such as “the BF of 1/50 proves the null hypothesis” is incorrect. Related to the previous comments regarding choosing a suitable prior distribution, if a prior distribution is not appropriate, the resulting BF is likely to be biased against it, thereby inaccurately estimating the strength of evidence.

## Conclusion

Relying on *p*-values alone may only provide a limited perspective of research findings. Bayesian analyses provide an alternative to traditional statistical analyses by enriching the information extracted from the data. The validity of research findings is the foundation of the scientific evidence that contributes to translational research and evidence-based practice. As demonstrated in this paper, Bayesian analyses are no more difficult to understand and interpret than traditional analyses. Researchers are encouraged to incorporate results of previous research into their current studies through the use of Bayesian statistics, thereby increasing the robustness of the results reported to inform future research and increase field knowledge.

## Declarations

### Ethics approval and consent to participate

Not applicable.

### Consent for publication

Not applicable.

### Availability of data and materials

The Nagy-Szakal et al. data is available at the Metabolomics Workbench repository under Project ID PR000576 (DOI: http://dx.doi.org/10.21228/M86X1F).

The Che et al. data is available at the Metabolomics Workbench repository under Project DOI 10.21228/M8PD9N

The Naviaux et al. data is available at the Metabolomics Workbench repository under Project DOI 10.21228/M82K58

### Authors’ contributions

WIL conceived and designed the ME-CFS studies. OF generated data. CB, OF conceived and designed the Bayesian studies. CB conducted statistical analyses and wrote first manuscript draft. XC performed statistical analyses in comparison to Bayesian studies. All authors edited and approved the manuscript.

## Acknowledgements

The authors acknowledge the contributions and participations of subjects living with ME/CFS.

